# Molecular characterization of pathogenic African trypanosomes in biting flies and camels in surra-endemic areas outside the tsetse fly belt in Kenya

**DOI:** 10.1101/2020.06.18.156869

**Authors:** Merid N. Getahun, Jandouwe Villinger, Joel L. Bargul, Abel Orone, John Ngiela, Peter O. Ahuya, Jackson M. Muema, Rajinder K. Saini, Baldwyn Torto, Daniel K. Masiga

## Abstract

**Background:** African animal trypanosomosis is becoming prevalent beyond its traditionally defined geographical boundaries and is a threat to animals beyond the tsetse belts in and outside Africa. However, knowledge of infections with clinically important trypanosome species and their diversity among field-collected hematophagous biting flies and domestic animals is limited mainly to tsetse and their mammalian hosts in tsetse-infested areas. This study aimed to examine the presence of trypanosomes in both biting flies and domestic animals outside the tsetse belt in northern Kenya, potential mechanical vector species, and their host-feeding profiles.

**Methods:** We screened for pathogenic African trypanosomes in blood samples from domestic animals and field-trapped flies by microscopy and sequencing of internal transcribed spacer (ITS1) gene PCR products. We sequenced kinetoplast maxicircle genes to confirm *Trypanosoma brucei* detection and the RoTat 1.2 and kinetoplast minicircle genes to differentiate type-A and type-B *Trypanosoma evansi*, respectively. Further, we identified the hosts that field-trapped flies fed on by PCR-HRM and sequencing of 16S rRNA genes.

**Results:** *Hippobosca camelina*, *Stomoxys calcitrans, Tabanus* spp., and *Pangonia rueppellii* are potential vectors of trypanosomes outside the tsetse belt in Marsabit County, northern Kenya. We identified *Trypanosoma spp*., including *Trypanosoma vivax*, *T. evansi*, *T. brucei*, and *T. congolense* in these biting flies as well as in camels (*Camelus dromedarius*). Trypanosomes detected varied from single up to three trypanosome species in *H. camelina* and camels in areas where no tsetse flies were trapped. Similar trypanosomes were detected in *Glossina pallidipes* collected from a tsetse-infested area in Shimba Hills, coastal Kenya, showing the wide geographic distribution of trypanosomes. Furthermore, we show that these biting flies acquired blood meals from camels, cattle, goats, and sheep. Phylogenetic analysis revealed diverse *Trypanosoma* spp. associated with variations in virulence and epidemiology in camels, which suggests that camel trypanosomosis may be due to mixed trypanosome infections rather than only surra (*T. evansi*), as previously thought.

## 1. Introduction

*Trypanosoma evansi* is one of the most important *Trypanosoma* spp. infecting livestock globally. Its wide geographic distribution [1–3], mode of transmission [4, 5], zoonotic potential [6, 7], pathogenicity to several domestic animals [8], high genetic diversity, and variation in virulence [5, 9] makes it an important parasite. Animal trypanosomosis caused by *T. evansi* is called surra in camels and is the most lethal disease of camels worldwide [4, 10, 11]. In addition to causing camel mortality, *T. evansi* infections reduce production of milk, an important staple food and the primary source of protein for pastoralists. Furthermore, *T. evansi* is an important pathogen in cattle and buffalo that results in morbidity and mortality and induces higher rates of abortion in domestic animals in Asia [11, 12].

Basic knowledge of the epidemiology and diversity of clinically important trypanosomes (such as *Trypanosoma vivax*, *T. congolense*, and *T. evansi*) in non-tsetse hematophagous flies and domestic animals from tsetse-free areas is significantly outweighed by that of tsetse and trypanosomes studied in tsetse-infested areas. However, several laboratory and semi-field experiments have demonstrated that different *Trypanosoma* spp., including *T. congolense* [13], *T. vivax* [14], and *T. evansi* [15], are potentially transmitted to domestic animals by various biting flies, such as *Stomoxys* spp. and *Tabanus* spp.

To evaluate the presence of various *Trypanosoma* spp. and the potential role of non-tsetse biting flies in their transmission within a tsetse-free area of northern Kenya, we investigated: (i) the diversity of hematophagous biting flies that could be involved in the mechanical transmission of trypanosomes; (ii) the identity, and diversity of economically and clinically important pathogenic trypanosomes within randomly collected and sampled flies and domestic animals, and (iii) on what vertebrates these biting flies feeding.

## 2. Materials and Methods

### 2.1. Study sites

Among three study sites, two sites in Marsabit County, northern Kenya, were in Ngurunit (N01°.74’, E 037.29’) at the edge of the tsetse distribution map and Shurr, a tsetse free area (N02°.08’, E038°.27’) (Fig. 1A-B). The third site was in Nanyuki in central Kenya, which is also tsetse free area (N00°.41’, E036°.90’) (Fig. 1A). All the three study sites are characterized by arid and semi-arid climatic conditions. The main means of livelihood is animal husbandry. The area has suitable biomass, especially for browsers such as goats and camels. Even though Ngurunit falls within the tsetse distribution map, a previous study reported no tsetse flies in the area [15] and therefore ideal sites to study non-tsetse transmitted trypanosomes. Furthermore, Marsabit County, where the two sampling sites are located have high number of camels, 1.3 – 1.9 camels/km^2^ [16], making it suitable for camel trypanosomosis studies. Tsetse flies (*G. pallidipes*) were collected from Shimba Hills in Kwale County, coastal Kenya for *Trypanosoma* spp comparisons.

**Fig. 1.**
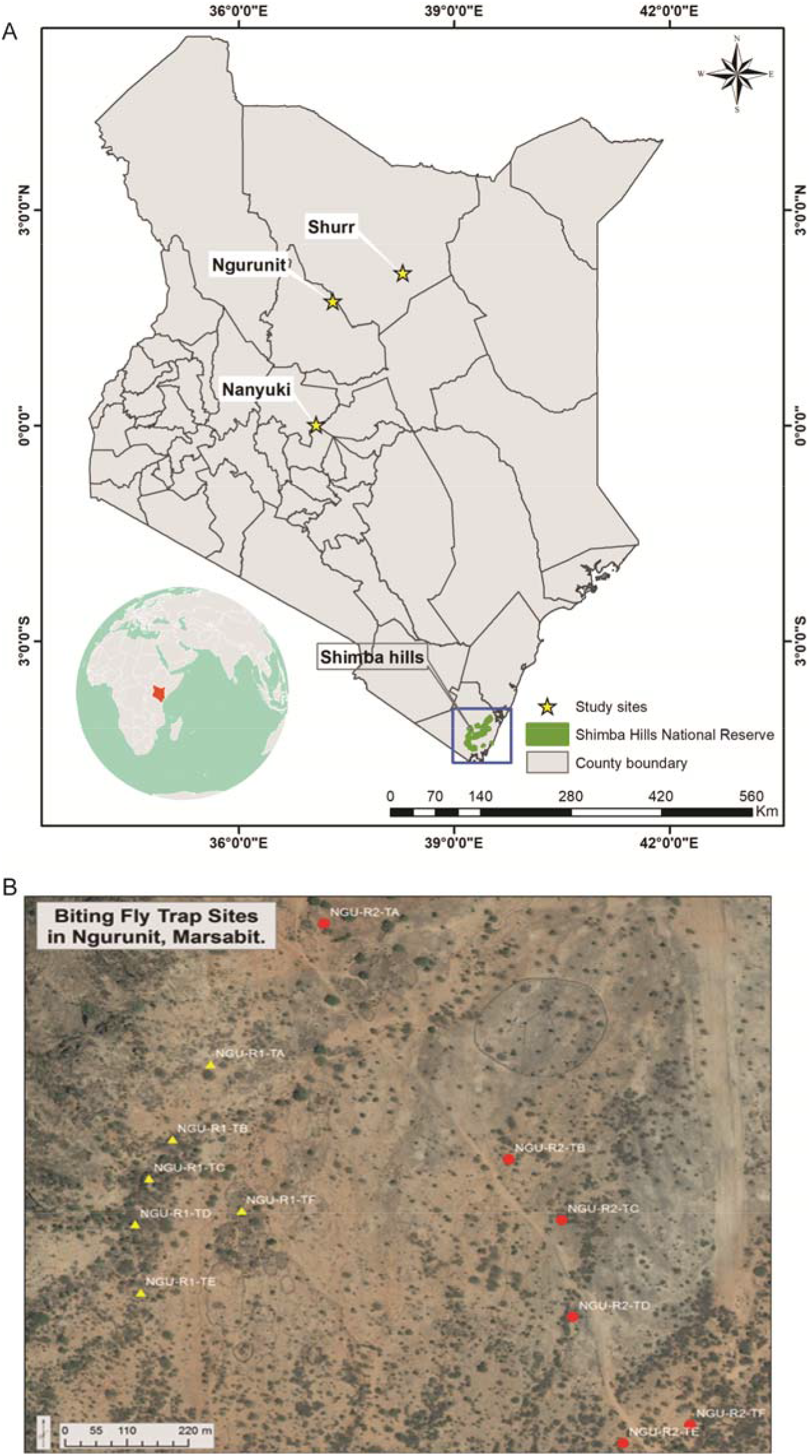
Map of study sites. (A) Map showing the study sites (B) Map of biting fly trap sites in Ngurunit village in Marsabit County. Yellow triangles represent the first replication of fly trapping in Ngurunit, which was in a forested area, and red circles represent the second replication, which was in an area with sparse vegetation.

### 1.2. Fly trapping

Flies were trapped using monoconical traps [17], placed ~150 m apart. Camel urine odour dispensed from plastic bottles (release rate was not quantified) was used as an attractant. Biting flies feeding on camels were collected using sweep nets and preserved in absolute ethanol and identified using appropriate taxonomic keys [18]. Camels, from which biting flies were collected, were randomly chosen regardless of their sex, age, or health status of the animal. The flies were trapped during both dry and rainy seasons using 25 monoconical traps per site for five days at each season and emptied every 24 hr. The fly trapping sites included thick bushland, *Acacia* spp. woodlands, watering points, animal enclosures, and open grassland areas (Fig.1B).

### 2.3. Fly density per camel

*Hippobosca camelina* and *Stomoxys calcitrans* counts were made by approaching ten randomly selected camels slowly from the side. An estimate number of *S. calcitrans* and *H. camelina* was made by counting the total number of flies on the legs and belly of the camel. Counts were made from a distance of 0.5 to 1 m. Two experienced technicians with expertise in distinguishing between the target flies performed all the counting. Stable flies could be differentiated from morphologically-similar house flies by their distinctive feeding posture as they feed parallel to the camel’s body [19], whilst house flies display a more random position when resting on the camel.

### 2.4. Blood sampling and microscopy

Approximately 5-10 mL of blood was drawn from the jugular vein of camels, goats, sheep, donkeys, and cattle into vacutainer tubes containing disodium salt of ethylene diamine tetra-acetate (EDTA) (Plymouth PLG, UK). An aliquot from each vacutainer tube was transferred into heparinized capillary tubes (75 × 1.5 mm) and spun in a micro-haematocrit centrifuge at 12,000 rpm for 5 minutes to separate the red and white blood cells and plasma, hence concentrating the trypanosomes [20]. Packed cell volume (PCV), an indicator of the animal’s anaemic status, was measured using Haematocrit Reader (Hawksley & Sons Limited, England) and expressed as a percentage of PCV to total blood volume. The buffy coat plasma interface was placed onto a microscope glass slide and examined under a microscope for the presence of moving trypanosomes. The trypanosome species were provisionally identified based on cell motility and morphology using wet blood film examination [20, 21]. Furthermore, thin blood smears were prepared from the samples, fixed with methanol, and stained with 10% Giemsa [20]. The stain was flushed with running tap water and allowed to dry for 35 minutes. Slides were then examined under the 100× oil immersion objective for trypanosomes and positive cases recorded. The rest of the well-mixed blood contents were appropriately labelled and stored in liquid nitrogen for transportation to Nairobi-based *icipe* laboratories for further screening. Domestic animals (including camels), were randomly sampled from various herds, for up to a maximum of 10% of the herd population to accommodate sampling of more herds. Camels in the herd were assigned a reference number and numbers were selected randomly for blood sampling. The mean number of camels per household varied from four to 35 on average in Ngurunit and Shurr sites, respectively. However, in Nanyuki, with relatively smaller population of camels, only one herd was sampled. All the identified infected camels were treated with triquin (Vetoquinol^®^) at a dose of 5 mg/kg body weight [10].

### 2.5 DNA extraction from blood and biting flies

Total genomic DNA was extracted from all collected blood samples using DNeasy Blood & Tissue Kits (Cat No./ID: 69504, Qiagen, Hilden, Germany), as follows: 100 μL of each blood sample was pipetted into 1.5-mL Eppendorf® tubes and mixed with 20 μL proteinase K, and the volume adjusted to 220 μL with PBS at pH 7.4. Subsequently, 200 μL Buffer AL were added to the reaction mix, thoroughly mixed and incubated for 10 minutes at 56°C. After the 10-minute incubation, 200 μL of absolute ethanol were separately pipetted into each tube and vortexed thoroughly before pipetting the mixture into the DNeasy Mini spin column placed in a 2-mL collection tube for centrifugation at 8000 rpm for 1 minute. The DNA samples on the spin columns were separately washed with 500 μL of Buffer AW1 Buffer AW2 preceding elution into clean 1.5-mL microcentrifuge tubes, with100 μL Buffer AE. The freshly eluted DNA samples were stored at −20°C until PCR analysis. Similarly, the total DNA from the crushed guts of biting flies were extracted and purified after brief surface sterilisation with ethanol and cleaning with distilled water. A negative extraction control was also performed for contamination assessment.

### 2.6 Identification of trypanosomes in the blood and fly samples

To check the presence of trypanosomes, a random subset of the flies from diverse biting flies collected were analysed from the whole fly using PCR. To check trypanosomes in blood we combined light microscopy with more sensitive molecular techniques, primarily using DNA-based markers [22] that enabled differentiation of trypanosome species and their subgroups. We employed PCRs targeting the internal transcribed spacer (ITS-1) gene fragment, which is a conserved gene across all African trypanosomes [23]. The diagnostic PCR assays were carried out in 10-μL reaction mixtures containing 5 μL 2× DreamTaq mix, 3 μL PCR water, 0.5 μL ITS-1 primers (F: 5’-CCGGAAGTTCACCGATATTG-3’; R: 5’-TTGCTGCGTTCTTCAACGAA-3’) [24] and 1 μL DNA template. PCR amplification conditions were programmed as follows: 95°C denaturation step for 1 minute, 35 cycles of 95°C for 30 seconds, 61°C for 30 seconds, 72°C for 1 minute and final extension of 72°C for 10 minutes. Additionally, primers designed to amplify kinetoplast 9S ribosomal RNA subunit (kDNA 12 (modified): 5′-TTAATGCTATTAGATGGGTGTGG-3′; kDNA 13: 5′-CTCTCTGGTTCTCTGGGAAATCAA-3′) [25] and CO1 (Tb_kDNA_COI_Max1: 5’-CCCTACAACAGCACCAAGT-3’; Tb_kDNA_COI_Max2: 5’-TTCACATGGGTTGATTATGG-3’) [26] genes were used to differentiate *T. brucei* from *T. evansi* as previously described. To separate *T. evansi* subtypes A and B, we used type A-specific primers targeting the VDG RoTat 1.2 gene (F: 5′-GCGGGGTGTTTAAAGCAATA-3′; R: 5′-ATTAGTGCTGCGTGTGTTCG-3′) and type B-specific primers targeting the minicircle gene (EVAB-1: 5′-ACAGTCCGAGAGATAGAG-3′; EVAB-2: 5′-CTGTACTCTACATCTACCTC-3′) [27, 28]. For each PCR, a negative PCR control (non-template control) was set up alongside samples (SF1). This enabled detection of contamination during PCR set up.

The PCR amplicons were purified using Quickclean II gel extraction kit (GeneScript USA Inc., Piscataway, USA) according to the manufacturer’s instructions. Briefly, DNA bands of interest were excised from the agarose gel with a sharp, clean scalpel (replaced for each sample) into sterile 1.5-mL Eppendorf tubes and three volumes of binding buffer II added to each sample. The reaction tubes were then incubated at 55°C for 10 minutes with occasional vortexing until all gels melted and a pale-yellow mixture was observed. The samples were then transferred into spin columns and centrifuged at 6000 ×g for 1 minute. Subsequently, the samples were washed with 650 μL wash buffer and centrifuged at 12000 ×g for 1 minute. The DNA samples were then eluted from the columns with 50 μL elution buffer into sterile 1.5-mL Eppendorf tubes and stored at −20°C. Confirmation of the purified DNA was performed by gel electrophoresis and sent for sequencing at Macrogen (Netherlands).

All nucleotide sequences were edited and aligned using the MAFFT plugin in Geneious software version 11.1.4 [29]. Sequence identities were revealed by querying in the GenBank nr database using the Basic Local Alignment Search Tool (www.ncbi.nlm.nih.gov/BLAST/). The aligned ITS-1 sequences were used to construct a maximum likelihood phylogenetic tree using PHYML v. 3.0 [30]. The phylogeny employed the Akaike information criterion [31] for automatic model selection and tree topologies were estimated using nearest neighbor interchange (NNI) improvements over 1,000 bootstrap replicates. The phylogenetic tree was visualized using FigTree v1.4.2 [32].

### 2.7. Bloodmeal source identification in biting flies by PCR – high-resolution melting (HRM) analysis

Genomic DNA from blood-fed *H. camelina* and *S. calcitrans* and known vertebrate whole blood samples as positive controls were isolated using DNeasy Blood and Tissue kit (Cat No./ID: 69504, Qiagen, Hilden, Germany). Fed flies were selected by observing engorged abdomens and then the guts of blood-fed flies were crushed and homogenized individually using ten 2-mm yttria-stabilised zirconia beads in 400 μL of cold homogenization phosphate buffered saline (PBS) to each tube on ice, then agitating for 10 seconds in a Mini-BeadBeater-16 (BioSpec, Bartlesville, OK, USA). The homogenates were then centrifuged for 10 seconds in a bench top centrifuge (Eppendorf, USA) at 1500 relative centrifugal force at 4°C. Aliquots of 210 μL of each homogenate were used for nucleic acid extraction.

High-resolution melt profiles from PCR amplicons of different vertebrate 16S rRNA DNA in 100 *H. camelina* were analysed in Applied Biosystems QuantStudio 3 real-time PCR system (Thermo Scientific, USA) and used to identify the various vertebrate bloodmeal sources in fed flies as previously described [33, 34]. Blood-meal profiles of vertebrates were specifically matched to selected domestic and wild animal positive controls. Control vertebrate host samples, including human, cow (*Bos taurus*), sheep (*Ovis aries*), warthog (*Phacochoerus africanus*), African buffalo (*Syncerus caffer*), goat (*Capra aegagrus hircus*), elephant (*Loxodonta africana*), Sprague Dawley rat (*Rattus norvegicus*), and camel (*Camelus dromedarius*) served as reference controls. The 10-μL PCRs reaction consisted of 1 μL DNA template, 6 μL of PCR water, and 2 μL of 5× HOT FIREpol EvaGreen HRM Mix (Solis BioDyne, Tartu, Estonia) and 0.5 μM concentrations of each primer (see [33, 34] for details). The PCR thermal cycling conditions for cytochrome b were set as follows: initial denaturation 95°C for 15 minutes, 35 cycles of denaturation at 95°C for 30 seconds, annealing 58°C for 20 seconds, extension 72°C for 30 seconds and final extension of 72°C for 7 minutes with final PCR products kept at 4°C. The annealing temperature of the 16S rRNA DNA marker was 56°C. Following PCR amplification, HRM analysis was performed within normalised temperature regions of between 65°C - 78°C and 88°C - 95°C. The different melt curve profiles of the samples were compared to the reference standards, and representative samples under each peak were selected for gene sequencing.

### 2.8 Data analysis

Biting fly densities on camels were compared using the Mann-Whitney test as the data was not normal following normality according to Levene’s test of homogeneity of variance. We used the following formula developed by [35]: n = ln(α)/ln(1−p) to determine the minimum number of camels and biting flies to be sampled for trypanosomes detection. We used sensitive molecular tools for pathogen detection, and we assumed that 3% of field collected flies and camels were infected at 95% confidence limit, thus, α = 0.05, P = 0.03 (probability of detecting infected biting flies, camel). Sample size n = ln(α)/ln(1−p) therefore n = −2.99/−0.03 = 99.6, which suggested that a minimum of 100 randomly sampled camels and biting flies were needed [35]. However, when biting flies were few, we analyzed 50 flies.

Chi-squared tests were used to compare differences in the number of trypanosomes detected in biting fly species, and among different domestic animals. The independent t-test was used to compare PCV values between infected and non-infected camels. Blood-meal sources were also compared using the chi-squared test. All analyses were performed using GraphPad software (GraphPad Software, Inc, USA). The Shannon diversity index (H) was used to characterise the diversity of pathogens in biting flies using percent prevalence data and the diversity index of biting flies between sites was analyzed using number of individuals per trap calculated using PAST 3.11 (www.folk.uio.no/ohammer/past/) [36]. The relative feeding index of *H. camelina* and *S. calcitrans* was calculated according to [36, 37] as follows; Wi = Oi/Pi, where, Wi = feeding ratio for livestock i, Oi = percentage of livestock, I, in the blood meals, Pi = proportion or percentage of livestock i available in the environment.

## 3. RESULTS

### 3.1. Diverse biting flies were identified as potential trypanosome vectors in the tsetse-free area

We collected a variety of biting flies of the Order Diptera from the study areas using monoconical traps (Table 1). Similar hematophagous flies were also observed feeding on camels, including *H. camelina* (Leach), *S. calcitrans* (L), *Pangonia rueppellii* (Jaenn), *Haematopota pluvialis*, and *Tabanus* spp. (Fig. 2A–F). *Hippobosca camelina* and *S. calcitrans* were present all year-round and observed feeding together on the same camel. The abundance of biting flies on a given camel varied between biting fly species. For instance, more *H. camelina* flies per camel (n = 10) were recorded as compared to *S. calcitrans* (*P* < 0.005 Mann-Whitney Test) (Fig. 2G). However, the number of other biting flies on camels (*P. rueppellii, Tabanus* spp., and *Hae. pluvialis*) were too low for meaningful comparisons.

**Fig. 2.**
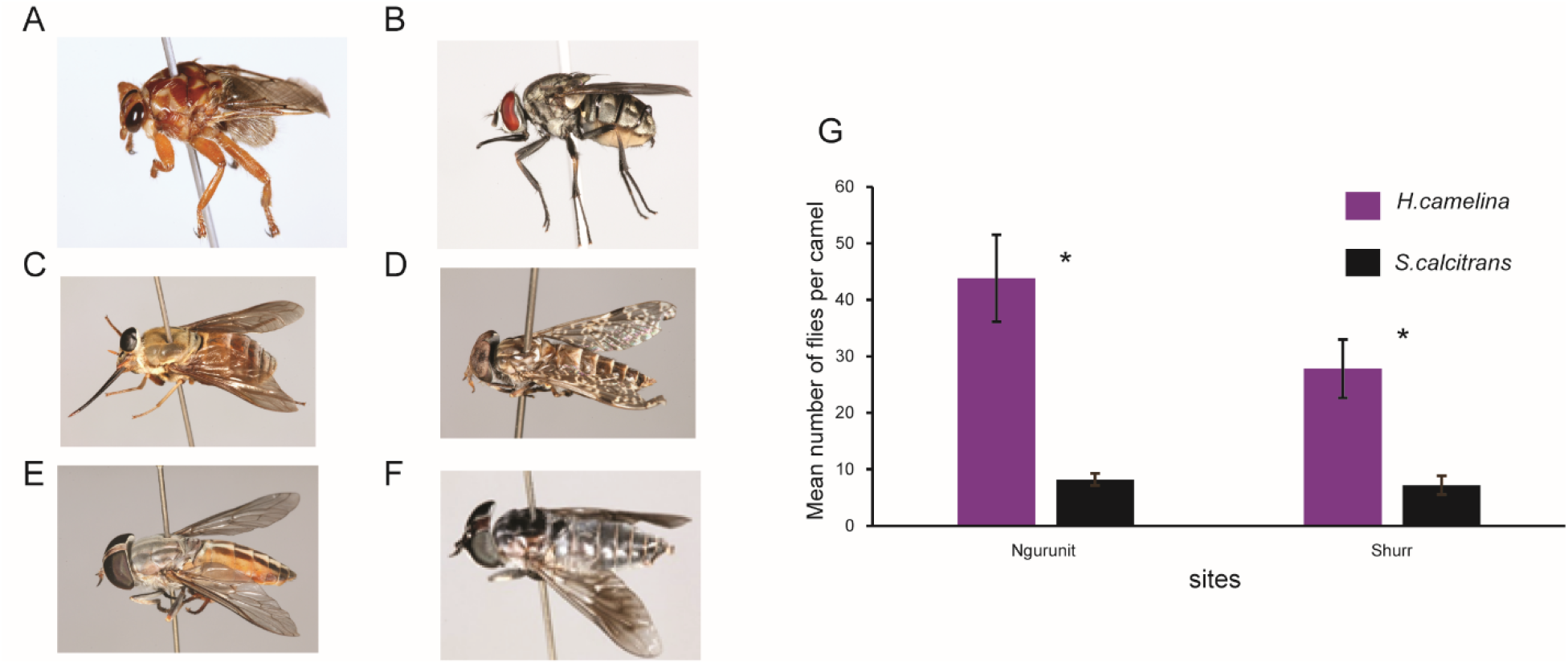
Diversity and abundance of biting flies. (A) *H. camelina*, (B) *S. calcitrans*, (C) *P. rueppellii*, (D) *Hae. pluvialis*, (E - F) *Tabanus* spp., (G) Mean number of *H. camelina* and *S. calcitrans* per camel at Ngurunit and Shurr sites, bars represent standard error of the mean, *depicts significant differences in fly density, *P* < 0.05, N = 10.

**Table 1.** Diversity of biting flies at three sites.

| Site | *H.*<br>*camelina* | *S.*<br>*calcitrans* | *Tabanus*<br>spp. | *P.*<br>*rueppellii* | *Hae.*<br>*pluvialis* | Shannon<br>index<br>(H) |
| --- | --- | --- | --- | --- | --- | --- |
| Ngurunit | + | + | + | + | + | 1.3 |
| Shurr | + | + | + | - | + | 0.6 |
| Nanyuki | - | + | - | - | - | 0 |

(+) indicates the specific biting fly was detected; (-) indicates not detected.

The diversity of biting flies varied from place to place; Ngurunit had diverse species of biting flies (Shannon index, *H* =1.3), including *H. camelina, P. rueppellii*, *Hae. pluvialis, S. calcitrans* and *Tabanus*. Shurr (Shannon index, *H* = 0.6) had all the species of biting flies, except *P. rueppellii*, while Nanyuki (Shannon index, *H* = 0) had only *S. calcitrans.*

### 3.2. *Trypanosoma* spp. identified in biting flies from the tsetse-free area detected using molecular technique

#### 3.2.1. Trypanosomes in *H. camelina*

Out of the 150 *H. camelina* (Fig. 2A) flies analysed, 3% had detectable *T. congolense* savannah PCR amplicons (700 bp), 39% had *T. vivax* amplicons (250 bp), 9% had *Trypanozoon* amplicons (500 bp), 6% had mixed *T. vivax* and *Trypanozoon* DNA, 3% had mixed *T. congolense* and *Trypanozoon* DNA, and 1% had DNA of all three species (*T. congolense*, *Trypanozoon*, and *T. vivax*) (Fig. 3A & B). *Trypanosoma vivax* was more common compared to *Trypanozoon* (χ^2^ = 36.883, df = 1, *P* < 0.001) and *Trypanozoon* was more common than *T. congolense* (χ^2^ = 4.771, *P* = 0.0289). Occurrence of mixed DNA was low (Fig. 3A–B). Trypanosomes were detected using total DNA isolated from individual flies by PCR targeting the trypanosomal ITS1 gene

**Fig. 3.**
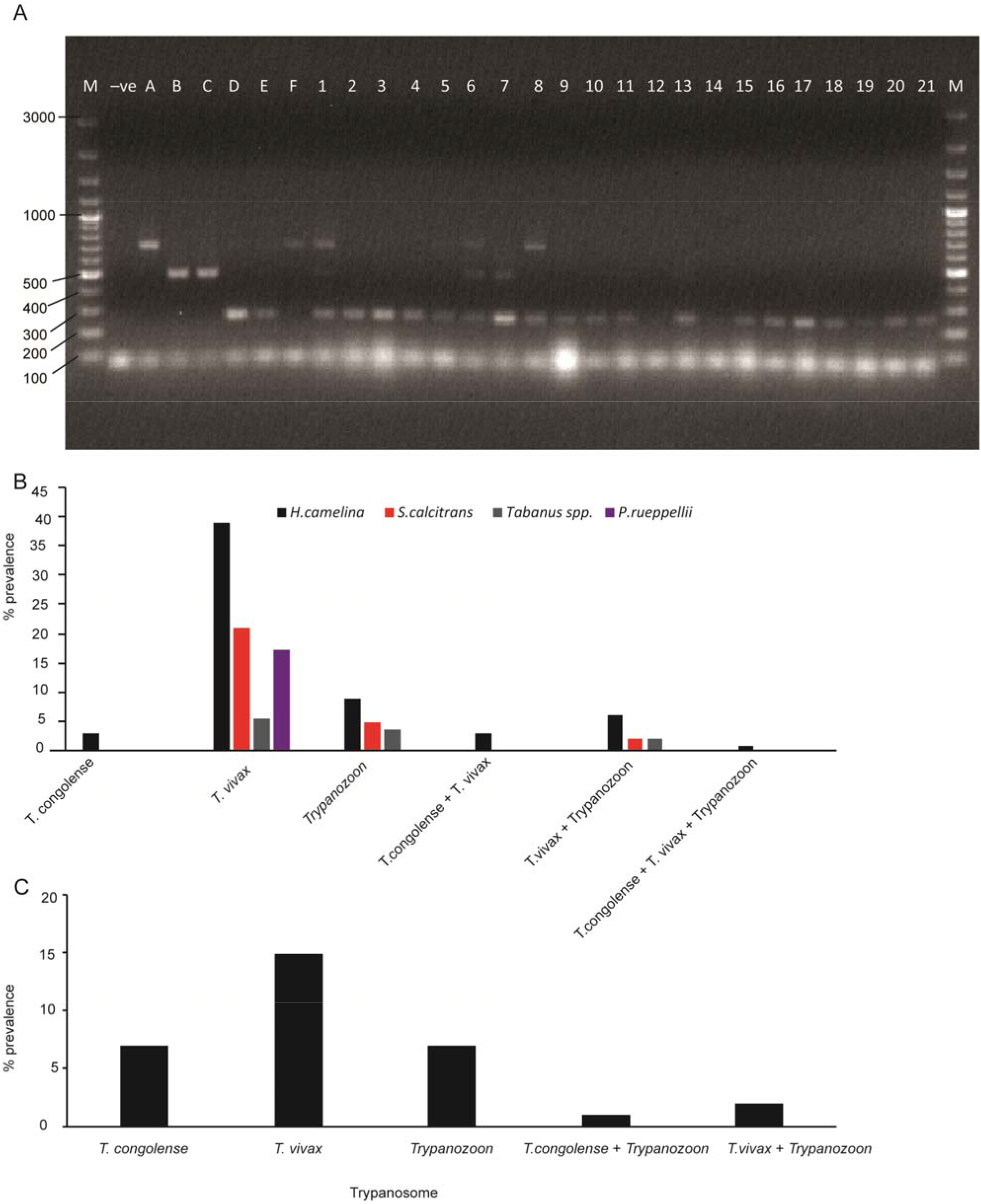
Prevalence and diversity of trypanosomes in biting flies. (A) PCR products were resolved through 1% ethidium-bromide stained agarose gel at 80V for 1.5 hrs. Lane M 100-bp marker (Thermo Scientific, USA); Lane ‘−ve’: negative control; Lane A – D contain positive controls for Lane A: *T. congolense* savannah (IL3000); Lane B: *T. brucei* ILTat 1.4; Lane C: *T. evansi* KETRI 2479; Lane D: *T. vivax* IL 2136; Lanes E and F: Trypanosome-infected camel blood from the same site as that of the flies; Lanes 1 – 21: some of selected *H. camelina* samples (**B**) The percentage prevalence of trypanosome species in biting flies. (**C**) The prevalence and diversity of trypanosomes detected from field collected *G. pallidipes* (n = 101).

##### 3.2.1.1 Diversity of trypanosomes in *H. camelina* from two sampling sites

The prevalence of trypanosomes in *H. camelina* collected from two sampling sites in northern Kenya, namely Ngurunit and Shurr, was variable. Out of the 150 flies analysed (75 flies from each site), 33% of flies from Ngurunit carried *T. vivax* (25/75), *Trypanozoon* was detected in 2.7% of flies (2/75), whereas about 5% of flies contained *T. congolense* savannah (4/75). However, 45% of *H. camelina* from Shurr were positive with *T. vivax* (34/75), whereas 14.7% of flies contained *Trypanozoon* (11/75). Thus, *H. camelina* collected from Shurr had significantly higher trypanosome-positive flies than those from Ngurunit (χ^2^ = 6.23, *P* = 0.013). However, *T. congolense* was not detected in flies sampled from Shurr flies. No *H. camelina* flies were found in Nanyuki (Table 1).

#### 3.2.2. Trypanosomes detected in *S. calcitrans*

Field-trapped *S. calcitrans* (n = 100), 50 from each of the two sampling sites in Marsabit (Fig. 2B) were analysed for the presence of trypanosomes; one fly from Ngurunit and four flies from Shurr were positive for *Trypanozoon* in total (5%) and in 21 flies *T. vivax* DNA was detected, eight from Ngurunit and 13 from Shurr site, in total, in 21% of flies. Similarly, more *S. calcitrans* were positive with *T. vivax* as compared to *Trypanozoon* (P < 0.001, χ^2^ test). Approximately 2% of the *S. calcitrans* from Shurr were positive for mixed *Trypanozoon* and *T. vivax* DNA (Figs. 3B and 4).

**Fig. 4.**
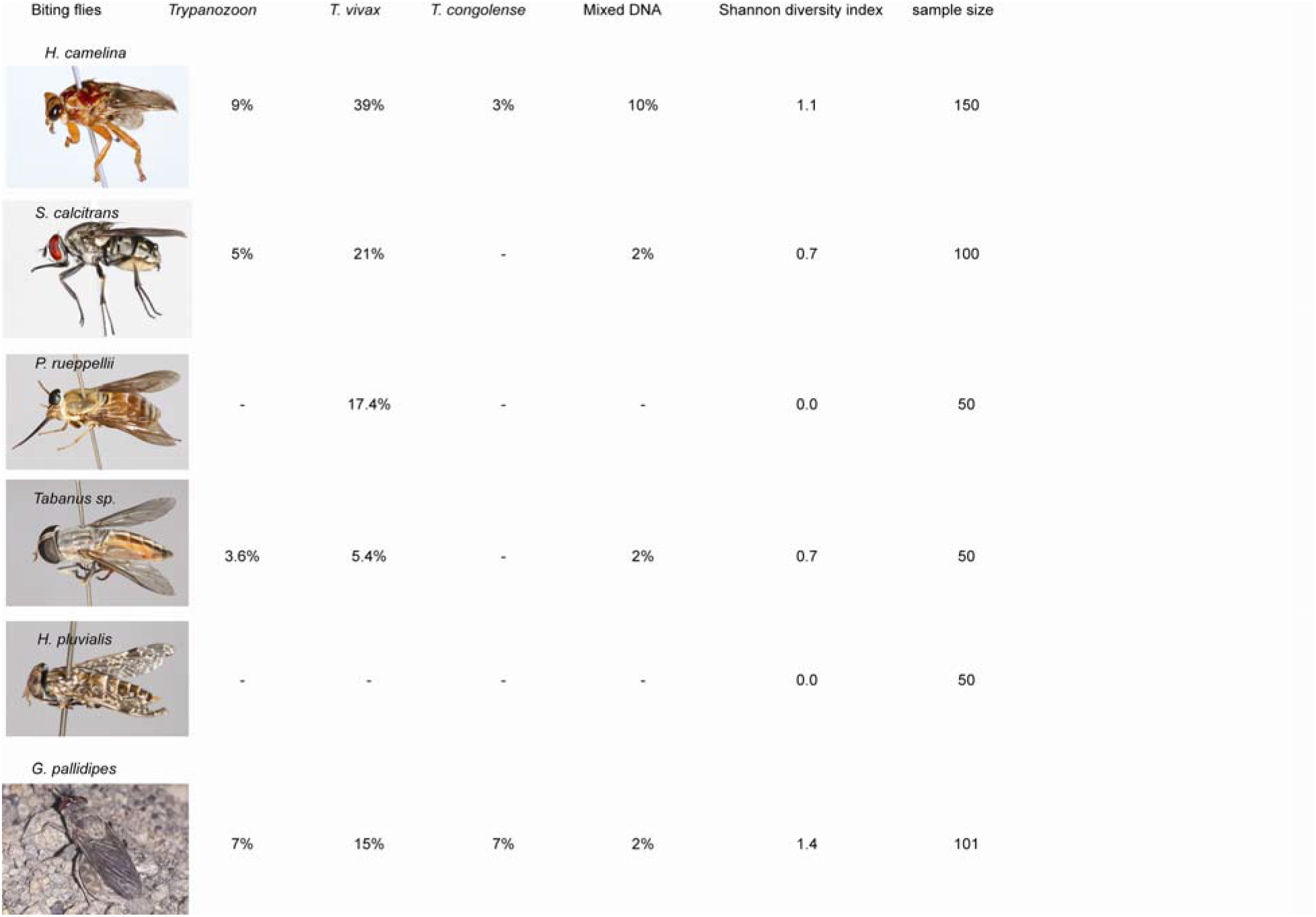
Diversity and percent of trypanosome species in biting flies. (−) indicates negative for that specific trypanosome species.

#### 3.2.3. Trypanosomes detected in *Tabanus* spp

Field trapped *Tabanus* spp. (Fig. 2E and F) contained *T. vivax* DNA (5.4%; n = 50 flies) and *Trypanozoon* (3.6%; n = 50 flies), trypanosomes DNA detected relatively lower in *Tabanus spp*. than in *H. camelina* and *S. calcitrans* (Figs. 3B and 4), and no significant difference was observed between the number of *T. vivax* and *Trypanozoon* DNA detected in the analysed *Tabanus spp.* flies (χ^2^, *P* > 0.05).

#### 3.2.4. Trypanosomes detected in *P. rueppellii*

We also identified another potential trypanosome vector, *P. rueppellii* (Fig. 2C) with *T. vivax* DNA in 17.4% of flies (n = 50), no other species of trypanosome was detected in *P. rueppellii* (Figs. 3B and 4). However, no trypanosomes were identified in 50 *Hae. pluvialis*, Fig. 2D. Among all biting flies analysed (except *Tabanus* spp.), *T. vivax* was the most abundant trypanosome species, followed by *Trypanozoon* (Fig. 3B).

#### 3.2.5. Trypanosomes detected in *G. pallidipes*

The trypanosomes diversity and prevalence were compared with freshly field-trapped trypanosome biological vector, *G. pallidipes*, from Shimba Hills in coastal Kenya (Fig. 1A). From 101 field-trapped *G. pallidipes* analysed, DNA of various trypanosomes were detected in 32%, 7.14% of which were *T. congolense* savannah (700 bp), 14.8% were *T. vivax* (250 bp), and 6.9% were *Trypanozoon* (500 bp). However, no significant differences in prevalence were recorded for the three trypanosome species, namely *T. congolense*, *T. vivax*, and *Trypanozoon* (either *T. brucei* or *T. evansi*) (*P* = 0.08). Similarly a low rates of mixed trypanosome DNA (*T.congolense* and *Trypaanozoon*) in 1% and in 2 % of *G. pallidipes*. *T. vivax* and *Trypanozoon* detected (Fig. 3C). Our results show that diverse non-tsetse hematophagous biting flies from the tsetse-free area harbour similar trypanosomes detected in *G. pallidipes* (Fig. 3C). The diversity of trypanosomes indicated by Shannon Index H varies between the different biting flies, *H. camelina* and *G. pallidipes* harboured more diverse trypanosomes DNA (Figure 4).

### 3.3. Trypanosome prevalence and diversity in camels

#### 3.3.1. Parasitological examination using light microscopy

Direct thin blood smears from the jugular vein showed an active infection of *Typanozoon* in 16 camels (7.2 ± 3.4%, 95% confidence intervals) of the sampled camels (n = 222). Trypanosomes were morphologically identified as belonging to *Trypanozoon* subgenus based on their morphology and motility. *T. vivax* is known to move fast but is less curved, whereas *T. brucei* moves fast but straight across a microscope field [21]. Trypanosome infections in camels varied in the parasite load from lowest parasitemia estimated at 10^6^ to highest of 5.10^8^ trypanosomes/mL blood (Figs 5A–B).

**Fig. 5.**
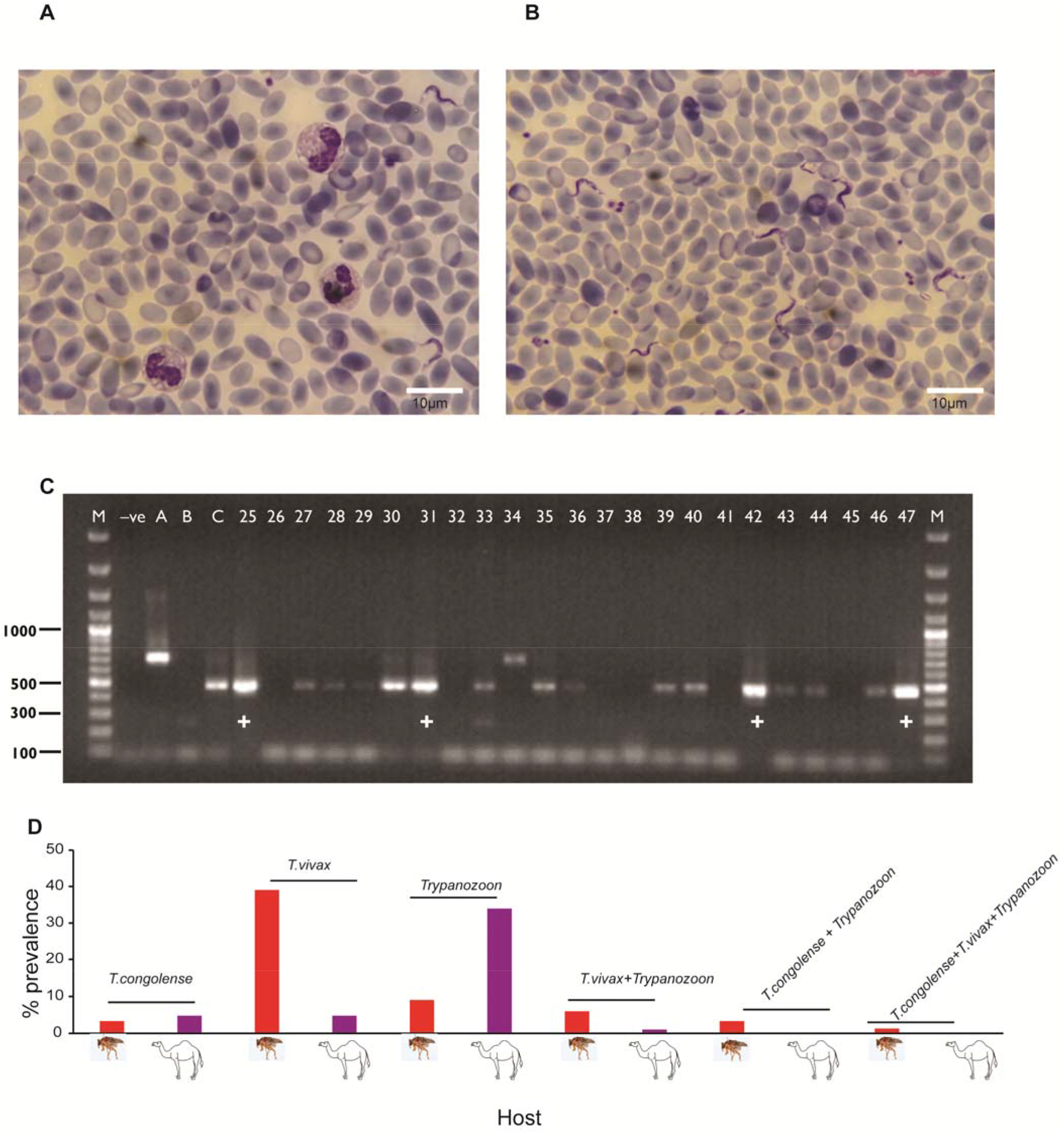
Detection of trypanosomes in camels and *H. camelina*. (**A - B**) Representative light micrographs of Giemsa-stained camel blood sample smears from two different camels showing varying parasitemia (A) low and (B) high parasitemia, *T. evansi* (GenBank accessions MH247174) with a small sub terminal kinetoplast at the pointed posterior end, a long free flagellum and a well-developed undulating membrane. (C) Agarose gel electrophoresis (1.2 %) performed on ITS1 PCR amplicons. Lane M: 100 bp marker; Lane ‘−ve’: negative control (non-template control); Lane A: *T. congolense* savannah IL3000; Lanes B: *T. vivax* IL2136; Lane C: *T. evansi* KETRI 2479; Lanes 25-47: selected camel blood samples, + shows camel that was trypanosome-positive by microscopy (D). Prevalence of trypanosome species in *H. camelina* and camels.

For further confirmation of the morphological identification, molecular analysis of microscopically positive and negative samples was performed. Figure 5C combines selected samples that were microscopically positive and those that were negative microscopically. Morphologically, *Trypanozoon*-positive samples 25 (GenBank accession MH247163), 31 (GenBank accession MH247168), 42 (GenBank accession MH247174), and 47 (GenBank accession MH247157) were also *Trypanozoon* positive by PCR and sequencing (Fig 5C) (for details, see section 3.3.2). In Fig 5C, six of the microscopically negative samples (26, 32, 37, 38, 41, and 45) remained negative with PCR. However, 13 samples (27-30; 33, 34-36; 39, 40, 43, 44, 46) that were negative microscopically, were found to be *Trypanozoon* positive with PCR, showing the sensitivity of PCR technique. A clear difference was observed in those microscopically positive and negative samples, relatively sharp and intense PCR band in those samples that were microscopically positive as compared to those microscopically negative (Fig 5C +). For further confirmation six microscopically *Trypanozoon* positive (GenBank accessions MH247157, MH247162, MH247163, MH247168, MH247170, and MH247174) and 10 microscopically negative, but *Trypanozoon* positive by PCR (GenBank accession MH247155, MH247158, MH247159, MH247160, MH247161, MH247166, MH247167, MH247169, MH247173 and MH247177) were sequenced (for details, see section 3.3.2).

#### 3.3.2. PCR-based trypanosome detection in camels

Trypanosomal ITS-1 PCR amplification showed 34% (75/222) of the camels were infected with *Trypanozoon*, which was more common than *T. vivax* infections (4.8%), with 250-bp PCR products (χ^2^ = 14.4, *P* < 0.001), and *T. congolense* savannah infections (4.8%), identified by 700-bp PCR amplicons. Mixed infections of *T. vivax* and *Trypanozoon* (*T. evansi* or/and *brucei*) accounted for 1.4% of the 222 camel samples analysed from the three sites. For further confirmation of size-based identification, we randomly selected 32 samples from PCR amplicons from each trypanosome species, those with good quality DNA and required concentration, for sequencing and those trypanosomes with known clinical symptoms. We sequenced 32 selected samples from all trypanosome species for further confirmation. The PCR amplicons of 32 samples were successfully sequenced and obtained sequences that clustered with known *T. vivax* or *T. evansi* and *T. brucei* isolates (Fig. 6). Five camels (GenBank accessions MH247140, MH247142, MH247145, MH247147, MH247149) and two *H. camelina* (GenBank accession MH247150, MH247152) were infected with *T. vivax* (100% nucleotide identity to GenBank accession MK880189). Two camels, one *H. camelina*, and one *S. calcitrans* were infected with trypanosomes (GenBank accessions MH247172, MH247174, MH247175, MH247177) sharing >99% ITS-1 nucleotide sequence identity with *T. evansi* sequences. Twenty-four sequences (GenBank accessions MH247139, MH247141, MH247146, MH247148, MH247151, MH247153-MH247163, MH247165-MH247170, MH247173, MH247176) clustered with *T. brucei*, sharing 99.3% identity with *T. brucei* (GenBank accession X05682), but also < 98% identity with *T. evansi* reference sequences. Further attempts to amplify and sequence *T. brucei-* specific kinetoplast maxicircle sequences, which are absent in *T. evansi*, confirmed four of the samples, one from *G. pallidipes* and three camel samples, as harbouring *T. brucei* (GenBank accessions MH247139, MH247159, MH247166, MH247167). We were unable to determine conclusively whether the other 17 samples with trypanosome ITS-1 sequences clustering among *T. brucei*, were *T. brucei* or *T. evansi* (Fig. 6).

**Figure 6:**
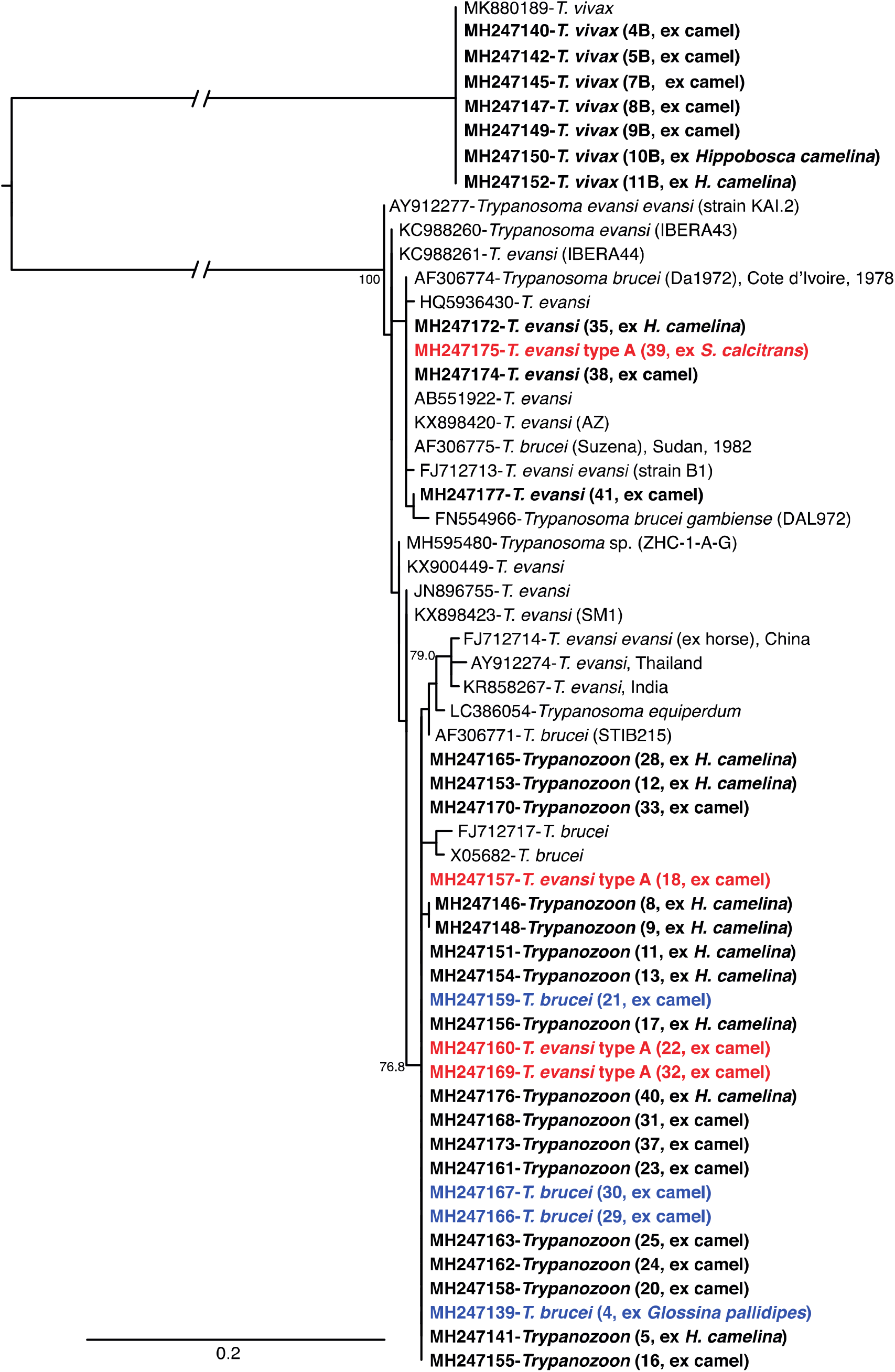
Maximum likelihood phylogeny of trypanosome ITS-1 nucleotide sequences. GenBank accession numbers and isolation sources are indicated. Sequences from this study are indicated in bold; trypanosomes were isolated from *G. pallidipes*, *S. calcitrans*, *H. camelina*, and camels. Sequences associated with samples confirmed to harbour *T. evansi* type A by amplification of the VDG RoTat 1.2 gene are highlighted in red. Sequences associated with samples confirmed to harbour *T. brucei* by maxicircle kDNA amplification are highlighted in blue. Bootstrap values at the major nodes represent agreement among 1000 replicates. The branch length scale represents substitutions per site. Branch gaps in the mid-point root branches represent 2.7 substitutions per site.

Further characterization of the samples using the *T. evansi* type A-specific primers differentiated the *T. evansi* in the *S. calcitrans* (GenBank accession MH247175) as *T. evansi* type A. Three of the camel samples with ITS-1 sequences (GenBank accessions MH247157; MH247160 and MH247169) that clustered among *T. brucei* also amplified using the *T. evansi* type A-specific primers, indicating either that *T. brucei* and *T. evansi* can share identical ITS-1 sequences or could be indicative of mixed infections with *T. brucei* and *T. evansi*. We failed to amplify *T. evansi* type B from *Trypanozoon* positive samples, though we cannot rule out the complete absence of Type B as we only analysed a subset of the samples.

#### 3.3.4 Anaemia associated with *Trypanozoon* infection

To determine the association between *Trypanozoon,* active infection and PCV, a measure of anaemia, which is one of the consequences of trypanosome infection [20, 39] and a symptom of surra, we compared the mean PCVs between camels with active *Trypanozoon* infection and those that were negative by microscopy. *Trypanozoon* infection affected the PCV values significantly; camels that had active *Trypanozoon* infection were anaemic with the mean PCV value of 24.64 ± 6; however, microscopically negative camels had a higher PCV value of 30.14 ± 6, t = 3.303, *P* = 0.002, n = 222).

### 3.4. Diversity of trypanosomes in other domestic animals co-herded with camels

We further analysed the prevalence of trypanosomes in other domestic animals such as goats, sheep, donkeys, and cattle that are co-herded with camels using PCR technique. *Trypanozoon* DNA was detected in small ruminants (goats and sheep), but not in cattle (Fig. 7). *Trypanosoma vivax* was more prevalent in sheep as compared to *Trypanozoon* (χ^2^ = 13.146, *P* = 0.0003, df = 1). In goats, *T. vivax* infections accounted for 8.2%, while the infection rate by *Trypanozoon* was 10.2% showing no significant difference (χ^2^, *P* > 0.05). Cattle were infected with *T. vivax* (9.5%) and *T. congolense* (2.4%) with no significant difference (Fig. 7). Furthermore, *T. vivax* was detected in 14% of donkeys, but no other trypanosomes species were detected (Fig. 7). However, all these livestock were negative microscopically.

**Fig. 7.**
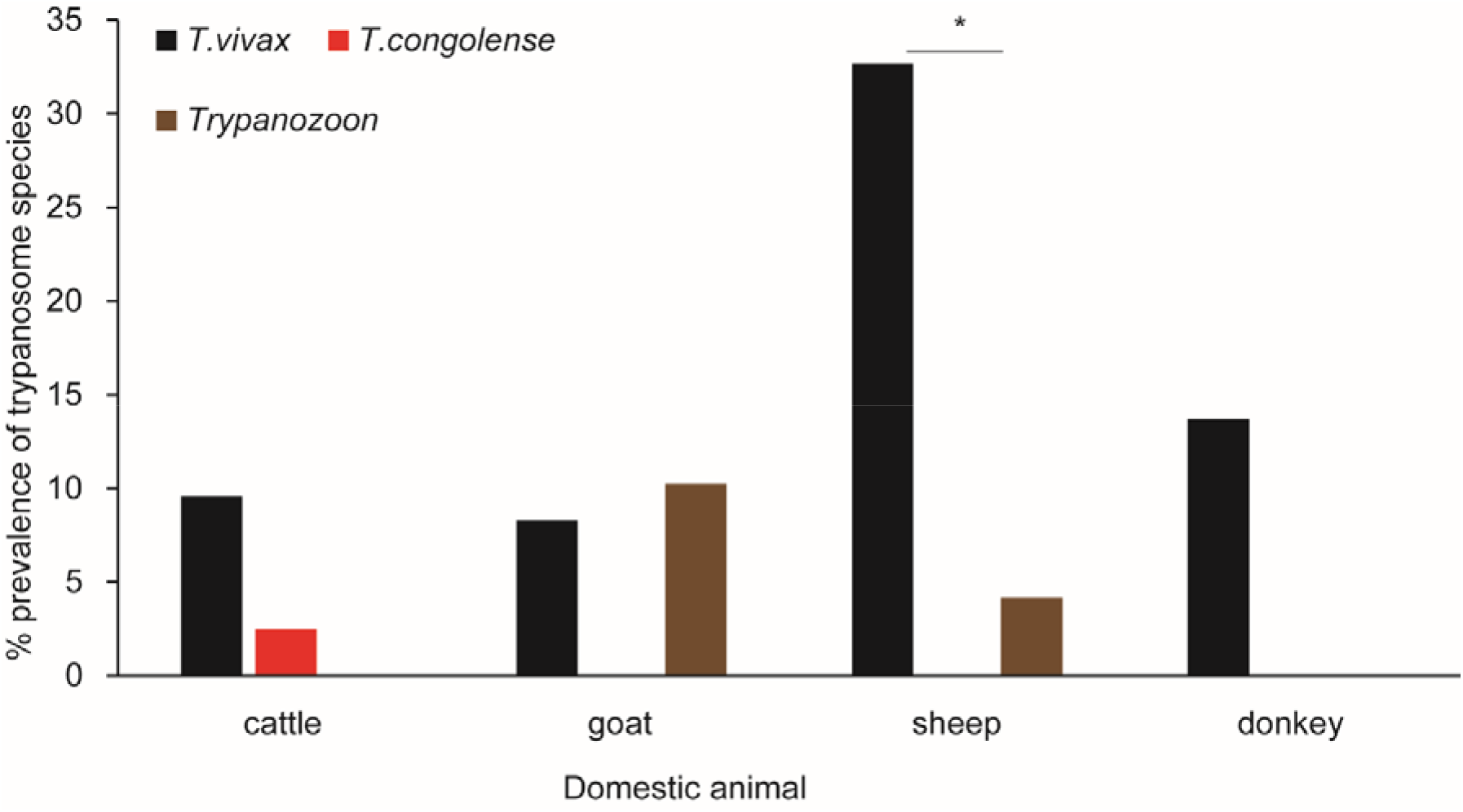
Prevalence and diversity of trypanosomes in domestic animals co-herded with camels. Percent prevalence of the three trypanosomes species in four domestic animals that co-herded with camels based on PCR. Significant difference in prevalence between trypanosomes species are depicted by an asterix. An equal number of blood samples were collected from Ngurunit and Shurr for cattle, goats, and sheep, while donkeys were exclusively sampled from Shurr.

### 3.5. Identification of vertebrate hosts from bloodmeal analysis of *H. camelina* and *S. calcitrans*

#### 3.5.1. Proportion of blood-meals taken by *H. camelina* from different hosts

The majority of *H. camelina* collected using traps had fed on domestic animals (95%, χ^2^ =, *P* < 0.0001). The camel host was the primary source of bloodmeals for 60% of the fed flies (χ^2^ = 7.96, *P* = 0.005), followed by goats (15%), sheep (14 %), and rats (4%) (Fig. 8A–B). Some of the flies fed on multiple vertebrate hosts; for instance, 1% of *H. camelina* fed on both human and sheep, 3% on camel and sheep, and 3% on goats and sheep. Each household kept diverse domestic animals (Fig. 8D). Sheep and goats were the most abundant, followed by camels, and relatively few cattle. The relative feeding index of *H. camelina* was calculated according to [37][38] as follow Wi = Oi/Pi, where, Wi = feeding ratio for livestock i, Oi = percentage of livestock i, in the blood meals, Pi = proportion or percentage of species i available in the environment. We calculated the relative feeding preference of *H. camelina* to camel against sheep, the most abundant livestock per household in Shurr. The average camel population per household was 35, while the average sheep population per household was 223. The camel: sheep abundance ratio was 35/223 = 0.157. 60% of *H. camelina* fed on camels, 15% fed on sheep, thus the observed feeding rate was 60/15 = 4. Therefore, the feeding index was 25.5 (4/0.157) as obtained from blood meal analysis of *H. camelina* shows a higher feeding preference on camels than sheep.

**Fig. 8.**
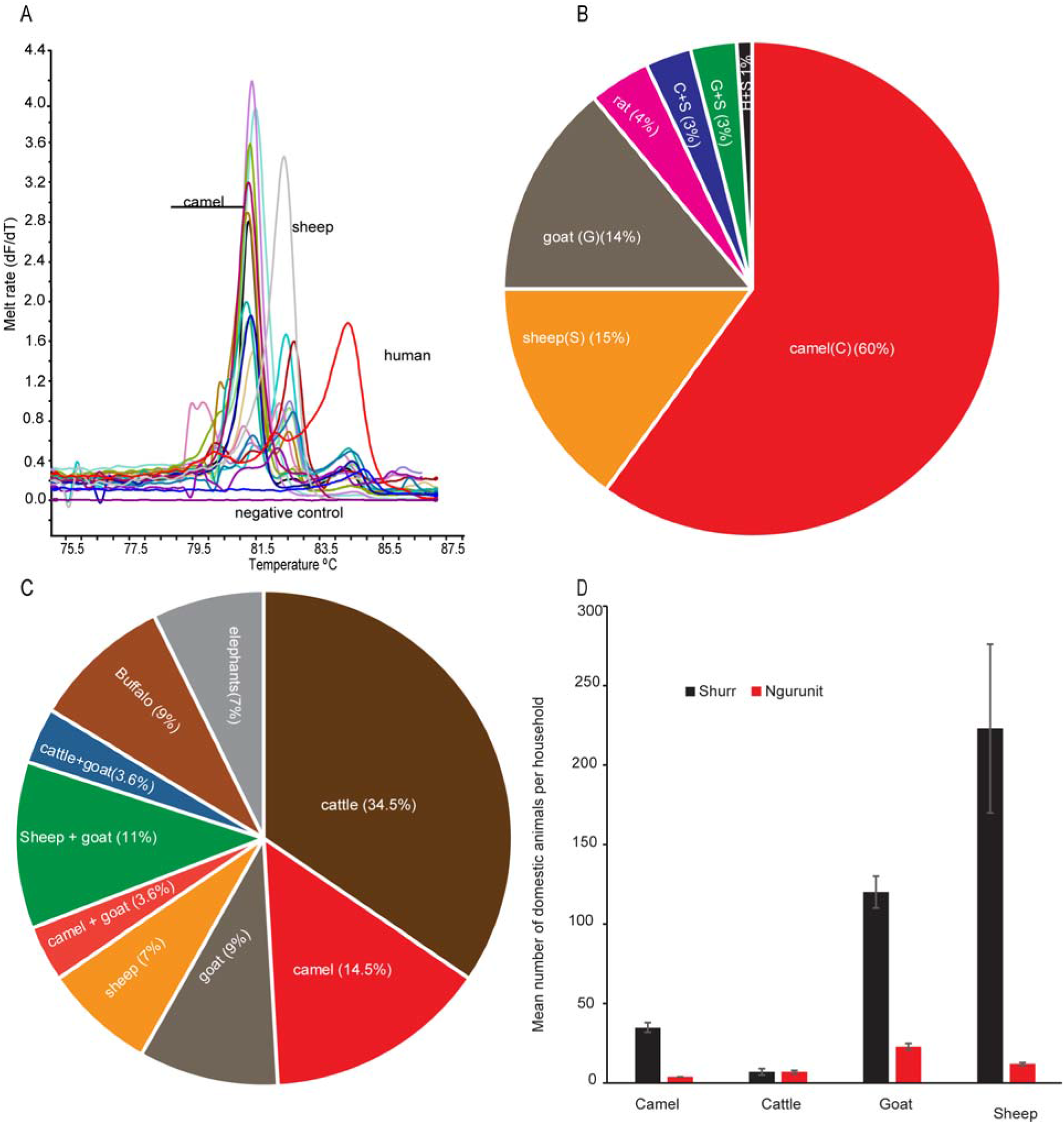
Bloodmeal analysis in *H. camelina* and *S. calcitrans* to determine vertebrate hosts. **(A)** Representative HRM profiles of selected *H. camelina* bloodmeal sources analyzed using 16S rRNA gene target. Positive controls were included for comparison. (**B**) Percent of bloodmeals for *H. camelina* identified using the 16S rRNA primer (n = 100). **(C)** Percent of blood meals from various vertebrate in *S. calcitrans* identified using16S rRNA primer (n = 40). **(D)** Diversity and abundance of domestic animals in Shurr and Ngurunit. Error bar indicates standard error, n = 96 and 92 households from Shurr and Ngurunit, respectively.

#### 3.5.2. Proportion of blood meals taken by *S. calcitrans* from vertebrate hosts

The bloodmeals of *S. calcitrans* included mainly cattle, camels, buffaloes, and goats (Fig. 8C). The relative feeding index of *S. calictrans* was calculated using the formula Wi = Oi/Pi described earlier in section 3.5.1. About 15% of *S. calictrans* fed on camel, 7% flies fed on sheep, observed feeding ratio 15/7= 2. Therefore, the feeding index of 12.7 (2/0.157) as obtained from blood meal analysis of *S. calcitrans* shows higher feeding preference to camel rather than sheep. However, *S. calcitrans* preferred to feed on cattle than camel. The camel: cattle abundance ratio was: 5 (35/7), and, the ratio of the number of feeds by *S. calcitrans* on camel and cattle, 0.428 (15/35). Thus, the feeding index for camel was 0.428/5 = 0.09, showing the high preference to feed on cattle than camel.

## 4. Discussion

Animal trypanosomosis is caused by several trypanosome species of genus *Trypanosoma* that are transmitted cyclically by tsetse flies and mechanically by other hematophagous flies. Since there were no tsetse flies in the study sites, other blood-sucking insects identified, such as *H. camelina* and stable flies, could serve as mechanical vectors that maintain trypanosomes in circulation among livestock. One of the factors that influences the mechanical transmission of trypanosomes by biting flies is the survival of the pathogen in the proboscis and midgut. Previous reports indicate that *T. evansi* can survive in the midgut of *H. camelina* for 75–90 minutes [15]. With our parasite establishment experiment *T. congolense* survived for 3 hrs and trypanozoon for 5 hr in the mid gut of *S. calcitrans*. Similarly, Sumba et al., 1998 [40] demonstrated that *T. congolense* survived for 3 and half hours and *T. evansi* for 8 hours in the gut of *Stomoxys niger* and *S. taeniatus*, showing their might be vectoral competence variation between *Stomoxys* species that need to be investigated. To determine how long it takes for *H. camelina* to find the next camel host when it is interrupted from feeding, we studied its mobility using a mark–release–recapture method by displacing them from the hosts. These flies were able to re-locate camels in ~60 minutes when translocated up to 1.5 Km away, which is within the time range of the parasite survival (SF2). Similarly, various *Stomoxys species,* which can transmit the pathogen at different time intervals and have long-distance mobility [13, 41].

The high number of *H. camelina* and *S. calcitrans,* their occurrence throughout the year, and the diversity of trypanosomes detected makes these two biting flies the most important potential vectors of trypanosome in tsetse-free ecologies. Furthermore, these two biting flies fed preferably on camels, but also on other hosts such as cattle and goats. However, the role of other biting flies identified, such as *Tabanus* spp. and *Pangonia* spp., should not be underestimated [13, 40, 42, 43]. These biting flies harbour similar trypanosome species as *G. pallidipes,* one of the major tsetse species in sub-Saharan Africa, due to its wider geographical distribution and economic importance [44–46]. The detection of diverse trypanosomes in non-tsetse infested areas shows that trypanosome diversity is conserved across a broader biogeography that includes tsetse-free regions.

The low percentage of the camels identified with active infection might be due to the low sensitivity of the microscopic technique [39, 47]. However, the molecular data showed that a significant number of camels were infected by trypanosomes, indicating that they might have subclinical infections. Thus, there is a possibility that sub-clinically infected camels can serve as a reservoir of trypanosomes. We also documented infected camels presenting with high parasitemia (Fig. 5B GenBank accession MH247157), which is required for mechanical transmission by biting flies [48]. Another reason for not detecting trypanosomes in blood might be due to trypanosomes moving to other tissues; for example, the skin is an anatomical reservoir of parasites of arthropod-borne diseases, such as trypanosomes [49]. Thus, we checked the lymph node aspirate (LNA) from a few sick camels with clear clinical signs of trypanosomosis, but with negative results during blood examination. We found that camel blood samples could test negative for trypanosome infection through microscopy and PCR but be positive in LNA (SF3 A-B). This finding agrees with previous reports that showed the presence of trypanosomes in LNA, but absence in the blood of cattle and sheep [50].

Camels that showed active infection microscopically were anaemic with average PCVs below 25, as compared to those microscopically negative camels with an average PCV of 30. With regards to different clinical outcomes, we have seen severely anaemic haemorrhagic camels that were infected by *T. vivax* (GenBank accessions MH247149) (PCV 9%) or by *T. evansi* type A (GenBank accessions MH247169) (PCV 11%). However, other camels with high *T. evansi* parasitaemia (millions of trypanosomes/mL of blood) were feeding well and in good body condition (PCV 21%) (GenBank accessions MH247174), which indicates that there could be different *T. evansi* populations as reflected in varying degrees of virulence to camels, that need further investigation. However, virulence also depends on the immune status of the host, past and recent infection by complex pathogens, as well as the genetic makeup of individual hosts. Similarly, *T. vivax-*infected camels with as low as 9% PCV have been reported previously[39]

We sequenced the trypanosome ITS-1 gene, which is the preferred target for species-specific molecular diagnostics of trypanosomes [23, 47]. The ITS-1 sequences produced distinct clusters for *T. vivax,* and *Trypanozoon* (Fig. 6). *Trypanosoma vivax* was the most prevalent trypanosome followed by *Trypanozoon*, both in flies and domestic animals, but not in camels, in which *Trypanozoon* was most common. We further detected maxicircle kDNA [25] in our *Trypanozoon* samples, confirming the presence of *T. brucei* in *G. pallidipes* and three camel samples. Additionally, we characterized *T. evansi* type A by the presence of the type A-specific marker targeting the RoTat 1.2 VSG gene, but we failed to detect *T. evansi* type B. The absence of *T. evansi* type B could be due to diagnostic limitations or the small sample size analysed, but these findings are congruent with previous findings of low occurrence of the *T. evansi* B subtype in camels [27, 28, 51]. However, for 20 of the *Trypanozoon* ITS-1 sequences obtained, we were not able to determine conclusively whether they were of *T. brucei* or *T. evansi,* showing that the ITS-1 gene cannot be used to effectively differentiate between these species. This challenge of *Trypanozoon* subgenus identification and the need for more specific and sensitive diagnostics is discussed by [52].

The detection of *T. brucei* and *T. congolense* both in biting flies and camel in the tsetse-free areas of northern Kenya might suggest the long-distance travelling of camels and other domestic animals between tsetse-free and the tsetse-infested areas, as some of the neighbouring counties are infested with tsetse flies. Similarly, previous studies detected *T. brucei* from camel in tsetse free area in northern Kenya [53][54]. To support this claim, a study done in North Eastern Kenya showed that animals move more than 120 Km from their homestead for better pasture and water, and infection of Rift Valley fever increases in herds that move than in those that remain at the homestead [55]. The other possibility is the presence of unidentified biological vectors, which will require further investigation, combined with vectorial capacity study of the identified biting flies and population genetics of trypanosomes.

## 5. Conclusions

The detection of diverse trypanosomes species/strains both in various biting flies and in camels suggests that trypanosomosis in camels is not only due to surra (*T. evansi* infection), but also nagana (*T. brucei*, *T. congolense*, and *T. vivax* infections). Such knowledge helps in drug administration because one could tailor the treatment to each trypanosome species [56]. Furthermore, our analysis shows that other domestic animals could serve as a reservoir of different trypanosomes, such as *T. vivax* and *T. congolense*, which are more deadly to camels [5, 9, 57]. Finally, the similarity of pathogens found in biting flies and their domestic animal blood-meal hosts demonstrates that these hematophagous flies could be used for xenomonitoring to track trypanosomes circulating in domestic animals as an early detection method.

## List of abbreviations

HRM: High-resolution melting
*icipe*: International Centre of Insect Physiology and Ecology
ITS1: Intergenic transcribed spacer gene 1
LNA: Lymph node aspirate
PCV: Packed Cell Volume
PCR: Polymerase chain reaction

## Declarations

## Acknowledgements

We thank Dr Robert Copeland for helping to identify the biting flies and taking the biting flies photo; Tiberius Marete, David Mbuvi, Peter Muasa and Irene Onyango for technical help; Kimathi, Emily and Barbara Kagima, for the study site map; Kelvin Muteru for his help in the bioinformatics; James Kabii for his technical help in the molecular studies, Collins Kigen, JohnMark, and Tawich K. Simon for technical help in the parasite survival experiments in *S. calcitrans* and Caroline Muya for handling the administrative issues. Special thanks go to Daud Tamasot (MCA), Malkash Lolkitarakino and Huka Guyo Qutte, Hussein Haji Abdulahi who facilitated our research at Shurr, Nanyuki and Ngurunit area. We thank Dr Mario Younan for his valuable technical advice on camels. We are grateful for the camel owners of Ngurunit, Nanyuki, and Shurr for their cooperation.

## Availability of data and materials

All data generated or analysed in this study are included in the article and as additional files. The newly generated sequences were deposited in the NCBI Nucleotide database under the accession numbers listed in Supplementary table.

## Authors’ contributions

MNG, BT, DM, SR conceptualized and designed the experiments. MNG, JLB, AO, POA, JN, JMM generated experimental data, JV contributed in the molecular part of the study MNG analysed the data and wrote the manuscript. All authors reviewed, edited and approved the final manuscript.

## Ethics approval and consent to participate

We collected blood samples within the framework of epidemiological surveillance activities, in accordance to the International Centre of Insect Physiology and Ecology*’s* Institutional Animal Care and Use Committee (IACUC) guidelines as performed during prophylaxis or diagnostic campaigns. Local authorities did not require ethical statements for the research studies. We did the blood sampling of domestic animals with the authorisation of the owner. Herdsmen/women gave their consent for their animal sampling after explaining the objectives of the study. No samples other than those for routine screening and diagnostic procedures were collected. All animals sampled and found positive with trypanosomes were treated using trypanocides.

## Consent for publication

Not applicable.

## Additional file

Additional file 1: Figure S1. PCR products were resolved 1% ethidium-bromide stained agarose gel (8V for 1.5 hrs) to check for any contamination. The DNA isolated from whole fly was amplified targeting trypanosomal ITS1 gene. Lane: M 10-bp marker, Bf (reaction buffer), wt (PCR water), TB (*T. brucei* ILTat 1.4) TV (*T. vivax* IL 2136), TC (*T. congolense* savannah (IL3000)), and TE (*T. evansi* KETRI 2479), F1-F10 DNA sample from *H. camelina* flies. The absence of PCR product under Bf, and wt show no contamination from extraction buffer.

Additional file 2: Figure S2, Number of *H.camelina* recaptured at the specificed distance from pint of release. Number in parentesis shows percentage of flies recaptured.

Additional file 3: Figure S3. (A) PCR products were resolved 1% ethidium-bromide stained agarose gel (8V for 1.5 hrs) to check for trypanosomes in blood and lymph node aspirate. The DNA isolated from blood and lymph node aspirate was amplified targeting trypanosomal ITS1 gene. Lane: M 10-bp marker, −Ve (reaction buffer), TE (*T. evansi* KETRI 2479) TV (*T. vivax* IL 2136), TC (*T. congolense* savannah (IL3000)), and LN_C1, LN_C2, DNA sample from two camels lymph node aspirate, B_C1 and B_C2 DNA from corresponding blood samples from the same camel. The result shows both samples of the lymph node aspirate were positive, while blood samples were negative from the same camel. (B) Five camels blood and lymph node aspirate were analysed, only camel five lymph node aspirate was positive for *T.vivax* but blood sample from the same camel was negative.

Additional File 2. Supplementary Table 1. Trypanosomes identified based on ITS1 gene sequence from different host included in Fig.6.

