## Supplementary Table 1 for "Molecular characterization of pathogenic African trypanosomes in biting flies and camels in surra-endemic areas outside the tsetse fly belt in Kenya"

Supplementary Table 1. Trypanosomes identified based on ITS1 gene sequence from different host included in Fig.6

| Sequence ID | organism | isolation-source | Host | Gene bank Accession number |
| --- | --- | --- | --- | --- |
| 4_ITS1 | *Trypanosoma brucei* | the whole fly | *Glossina pallidipes* | **MH247139** |
| 4B_ITS1 | *Trypanosoma vivax* | blood | camel | **MH247140** |
| 5_ITS1 | *Trypanozoon sp.* | the whole fly | *Hippobosca camelina* | **MH247141** |
| 5B_ITS1 | *Trypanosoma vivax* | the whole fly | *Hippobosca camelina* | **MH247142** |
| 7B_ITS1 | *Trypanosoma vivax* | blood | camel | **MH247145** |
| 8_ITS1 | *Trypanozoon sp.* | the whole fly | *Hippobosca camelina* | **MH247146** |
| 8B_ITS1 | *Trypanosoma vivax* | blood | Camel | **MH247147** |
| 9_ITS1 | *Trypanozoon sp.* | the whole fly | *Hippobosca camelina* | **MH247148** |
| 9B_ITS1 | *Trypanosoma vivax* | blood | Camel | **MH247149** |
| 10b_ITS1 | *Trypanosoma vivax* | the whole fly | *Hippobosca camelina* | **MH247150** |
| 11_ITS1 | *Trypanozoon sp.* | the whole fly | *Hippobosca camelina* | **MH247151** |
| 11B_ITS1 | *Trypanosoma vivax* | the whole fly | *Hippobosca camelina* | **MH247152** |
| 12_ITS1 | *Trypanozoon sp.* | the whole fly | *Hippobosca camelina* | **MH247153** |
| 13_ITS1 | *Trypanozoon sp.* | the whole fly | *Hippobosca camelina* | **MH247154** |
| 16_ITS1 | *Trypanozoon sp.* | blood | Camel | **MH247155** |
| 17_ITS1 | *Trypanozoon sp.* | the whole fly | *Hippobosca camelina* | **MH247156** |
| 18_ITS1b | *Trypanosoma evansi_A* | blood | Camel | **MH247157** |
| 20_ITS1 | *Trypanozoon sp.* | blood | Camel | **MH247158** |
| 21_ITS1b | *Trypanosoma evansi* | blood | Camel | **MH247159** |
| 22_ITS1 | *Trypanosoma brucei* | blood | Camel | **MH247160** |
| 23_ITS1 | *Trypanozoon sp.* | blood | Camel | **MH247161** |
| 24_ITS1 | *Trypanozoon sp.* | blood | Camel | **MH247162** |
| 25_ITS1 | *Trypanozoon sp.* | blood | Camel | **MH247163** |
| 28_ITS1 | *Trypanozoon sp.* | The whole fly | *Hippobosca camelina* | **MH247165** |
| 29_ITS1 | *Trypanosoma brucei* | blood | camel | **MH247166** |
| 30_ITS1b | *Trypanosoma brucei* | blood | camel | **MH247167** |
| 31_ITS1 | *Trypanozoon sp.* | blood | camel | **MH247168** |
| 32_ITS1 | *Trypanosoma evansi_A* | blood | camel | **MH247169** |
| 33_ITS1 | *Trypanozoon sp.* | blood | camel | **MH247170** |
| 35_ITS1 | *Trypanosoma evansi* | the whole fly | *Hippobosca camelina* | **MH247172** |
| 37_ITS1b | *Trypanozoon sp* | blood | camel | **MH247173** |
| 38_ITS1 | *Trypanosoma evansi* | blood | camel | **MH247174** |
| 39_ITS1 | *Trypanosoma evansi_A* | the whole fly | *Stomoxys calcitrans* | **MH247175** |
| 40_ITS1 | *Trypanozoon sp.* | the whole fly | *Hippobosca camelina* | **MH247176** |
| 41_ITS1 | *Trypanosoma evansi* | blood | camel | **MH247177** |
