## Supplementary figures and images for "Molecular characterization of pathogenic African trypanosomes in biting flies and camels in surra-endemic areas outside the tsetse fly belt in Kenya"

### Supplentary Fig 1

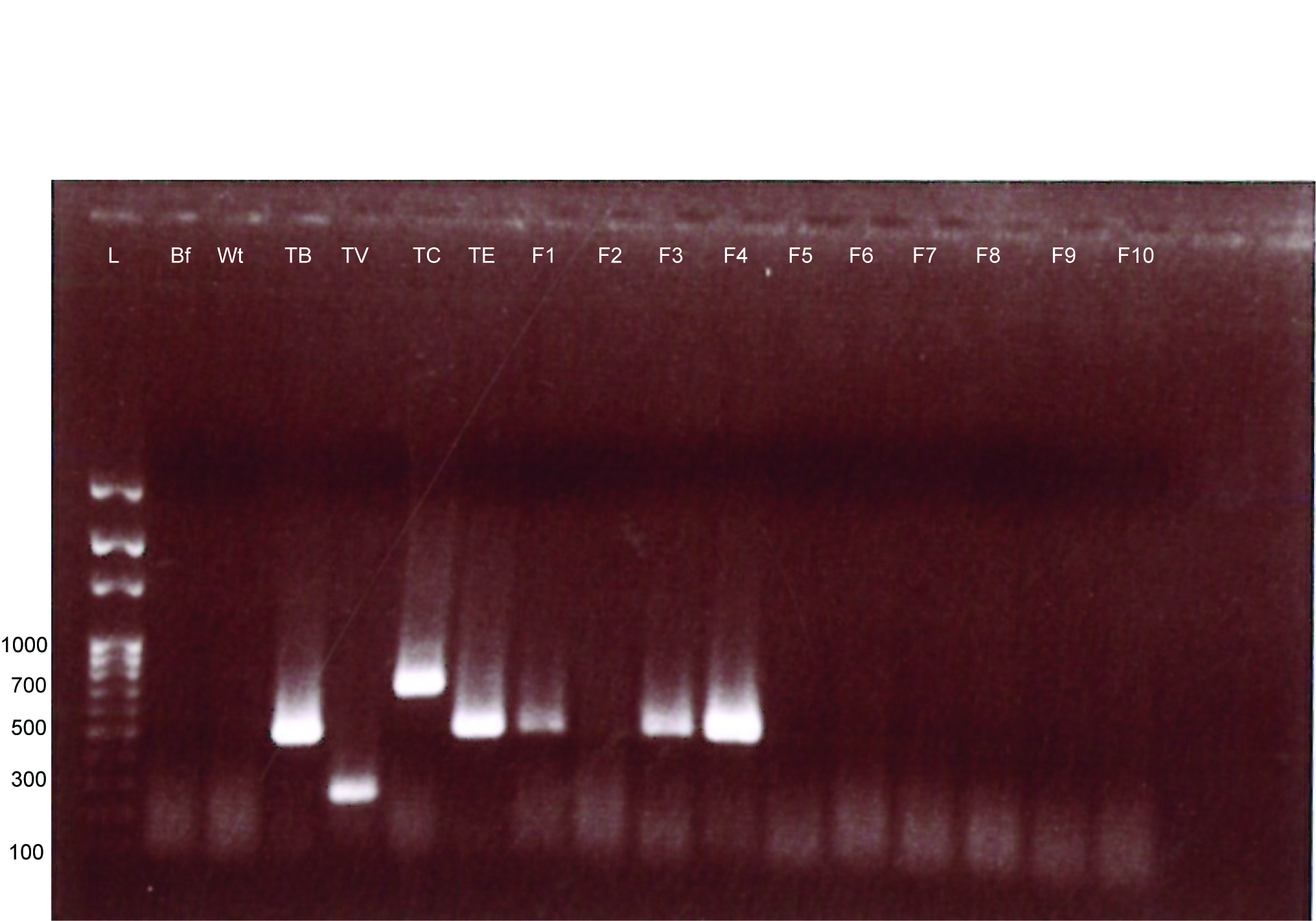

### Supplentary Fig 2

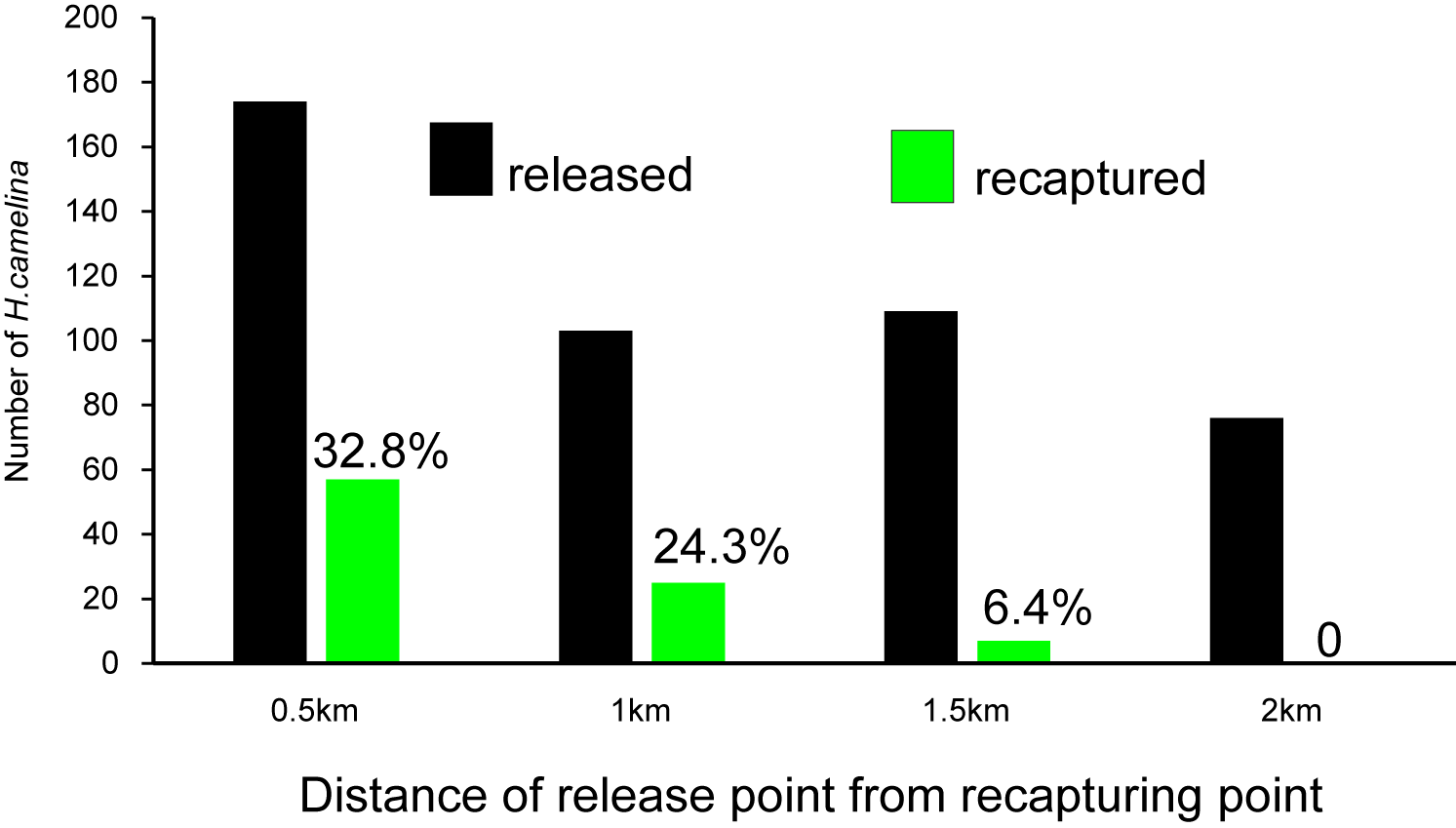

### Supplentary Fig 3

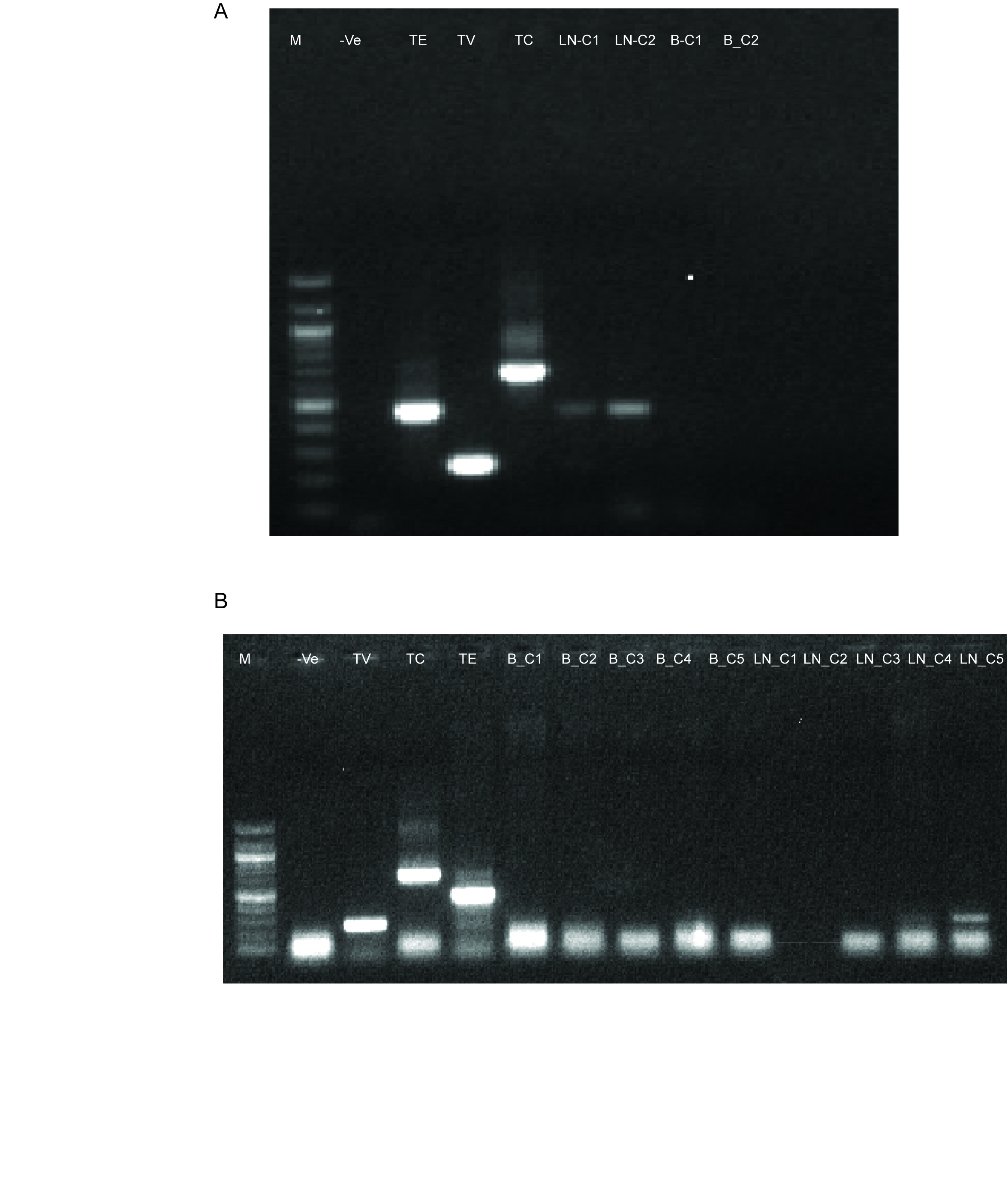
